# TASK-3 Two-Pore Potassium Channels drive neuronal excitability of the circadian clock and entrainment to challenging light environments

**DOI:** 10.1101/2020.07.27.222885

**Authors:** Aiste Steponenaite, Tatjana Lalic, Lynsey Atkinson, Liting Wei, Alistair Mathie, M. Zameel Cader, Gurprit S. Lall

## Abstract

The suprachiasmatic nucleus (SCN), the mammalian circadian clock, is a heterogenous structure made of several neuron types that generate a circadian electrical activity profile. However, it is unclear how such regulation in endogenous neuronal excitability is maintained. Background two-pore domain potassium channels (K2P), such as TASK-3, play an important role in inhibiting neuronal activity. Here, we utilize a TASK-3 KO mouse model to unravel the role played by this channel in SCN circadian neuronal regulation and behavioral photoentrainment. Our results reveal that TASK-3 is needed to adapt to challenging lighting conditions, such as those experienced through seasonal changes and jet lag. From our investigations this appears to be very distinct from pathways that drive acute, ‘one-off’ adjustments in clock phase, in response to single pulses of light. These findings provide crucial information on the intricate pathways linking clock output to behavioral adaptation to light-dark cycles.

## Introduction

Neurons of the suprachiasmatic nucleus (SCN) function as a molecular clock of the body orchestrating our daily lives and enabling adaptations to the environmental rhythms. This autonomous molecular clock works by the control of three transcription translation feedback loops (TTFL) with CLOCK and BMAL1 proteins being the initiators of the cascade of transcriptional events occurring at the nucleus (Ki *et al.*, 2015)(Patton and Hastings, 2018). Molecular clock proteins PER1 and PER2 induced upon light stimulation adjust the TTFL to the light/dark cycle and facilitate phase shifting responses happening during the subjective night as well as clock advances or delays based on the environmental light/dark cycle (Jagannath *et al.*, 2013)(Welsh, Takahashi and Kay, 2010).

Tight molecular clock regulation is also maintained by circadian variation of SCN neuron activity. Despite neuronal heterogeneity (Abrahamson and Moore, 2001), there is a clear pattern of electrical activity where SCN neurons endogenously generate high frequency action potentials during the day, which are silenced during the night. K^+^ dependent conductance hyperpolarizes membranes, hence rendering cells electrically inactive through altering the excitability threshold. Daily depolarization is mainly maintained by voltage gated Na^+^ and Ca^2+^ channels together with hyperpolarization activated cyclic nucleotide gated channels and NALCN sodium leak channel, bringing the membranes to more depolarized state and placing them near the threshold for generating an action potential (Patton and Hastings, 2018)(Allen, Nitabach and Colwell, 2017). At night hyperpolarizing K^+^ dependent conductance hyperpolarizes membranes in this way electrically inactivating the neurons with contribution by the BK channels.

Background two-pore domain potassium channels (K2P) play a major role in hyperpolarization by providing a leak K+ current over a wide voltage range (Feliciangeli *et al.*, 2015; Renigunta, Schlichthorl and Daut, 2015). There are 15 K2P family members, which are expressed throughout the central nervous system, including the autonomous circadian clock, suprachiasmatic nucleus (SCN). Regulation of the SCN is maintained by the rhythmic variation of its neuronal activity. Despite several studies investigating the actions of TASK-3 (Brickley *et al.*, 2007; Linden *et al.*, 2007), diseases and conditions linked to its mutations (Mu *et al.*, 2003; Pei *et al.*, 2003; Barel *et al.*, 2008; Williams, Bateman and O’Kelly, 2013; Sun *et al.*, 2016; Yoshida *et al.*, 2018; Liao *et al.*, 2019; Cooper *et al.*, 2020), the role of TASK-3 in the SCN is less well established. In this present study, we define the role of TASK-3 in the SCN through assessing its contribution in regulating and maintaining stable cellular and behavioral entrainment. We employ a TASK-3 knockout (KO) animal model to test the impact of TASK-3 on circadian regulation of cellular excitability and photoentrainment. Our findings show that TASK-3 is an essential component in governing both daily electrical output of the clock and robust light entrainment, driving environmental adaptation and survival in most species.

## Results

### TASK-3 expression is required for diurnal variation in resting membrane potential in SCN neurons

To establish a gene expression profile for TASK-3, SCNs from WT mice were collected every 4 hours (Fig. 1A). RT-qPCR shows that the expression of TASK-3 gene *Kcnk9* peaks during the late night, around ZT18 (6 hours following lights off). Following the observation of a diurnal regulation of *Kcnk9* mRNA levels in the SCN, we next sought to understand the impact on the electrical properties of the SCN. Examining the resting membrane potential of SCN neurons, we found WT SCN cells displayed a diurnal variation in membrane potential, with RMP at −48 mV during the day and −56.9 mV during the night (p=0.002, two-way ANOVA with Tukey’s multiple comparisons test). However, in the absence of TASK-3, this diurnal variation was abolished, and the SCN RMP resided at around −50mV regardless of time of day (p=0.999, two-way ANOVA, Tukey’s multiple comparisons test; Figure 1B). Two-way ANOVA showed a significant variance between genotypes (F_(1, 57)_ = 6.5, p=0.01), time of the day (F_(1, 57)_ = 7,7, p=0.007) and overall interaction (F_(1, 57)_ = 8.6, p=0.005). We have previously found similar results in TRESK KO mice (Lalic *et al*., in press).

**Figure 1.**
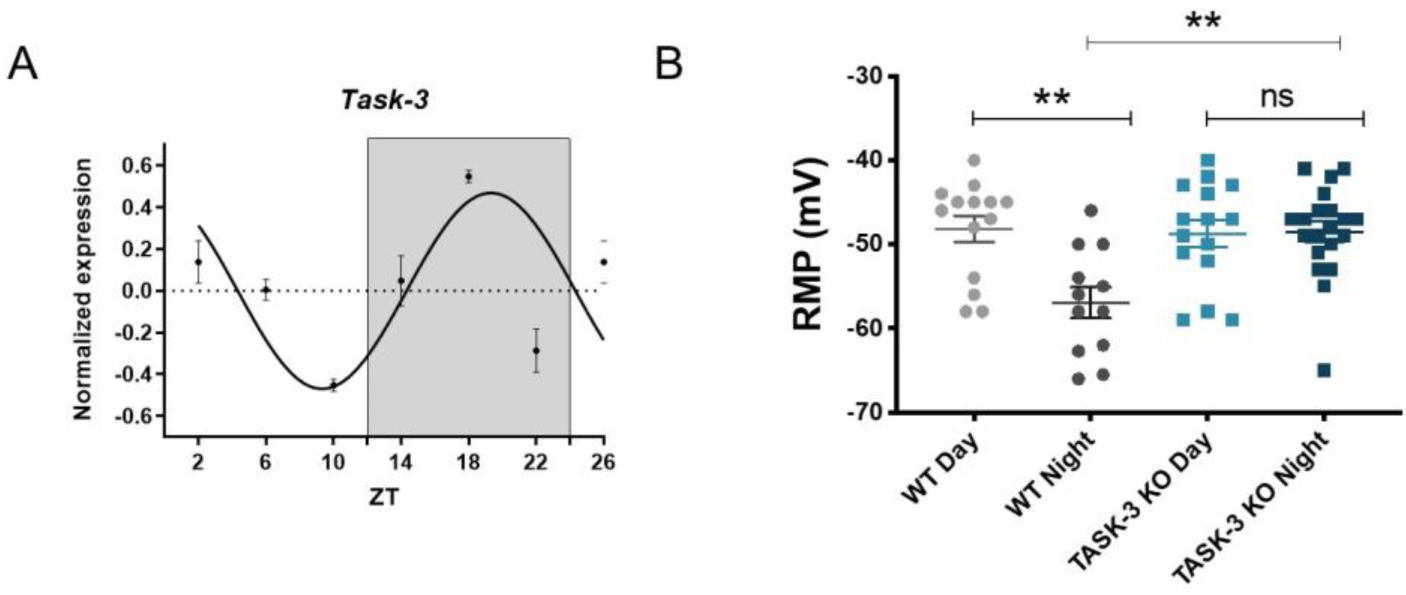
Task-3 diurnal variation in gene expression and RMP. (A) Circadian Task-3 mRNA expression in mouse SCN using RT-qPCR shows a clear diurnal profile peaking in the middle of subjective night (n=3). (B) Whole-cell, current clamp recordings of SCN neurons during the subjective day/night from WT and TASK-3 KO mice. TASK-3 KO mice lack diurnal variation in RMP, remaining in a constantly depolarized state. (WT: day, n=14; night, n=12; TASK-3 KO: day, n =15; night n=20; mean ± SEM).

Loss of TASK-3 results in increased nocturnal locomotor activity but preserved circadian clock resetting and diurnal behavior. WT and TASK-3 KO animals housed under a 12:12 light dark cycle showed stable synchronized activity profiles. However, TASK-3 KO animals exhibited elevated activity during the later night (Fig. 2A). Two-way ANOVA showed significance between genotypes (F_(1,576)_=37.7, p<0.0001), time of the day (F_(47,576)_=26.3, p<0.0001) and interaction (F_(47,576)_=2.4, p<0.0001). The overall 24hr infrared beam breaks were significantly elevated in the knockout mice; in line with previously published work (Fig. 2B, p<0.01, Student’s t-test). Housing animals under constant conditions of darkness showed free-running rhythms to be maintained in both WT and TASK-3 KO animals (Fig. 2C). However, the differences in overall activity remained under such environmental conditions. Two-way ANOVA showed significant variation between genotypes (F_(1,528)_=46, p<0.0001), time of the day (F_(43,528)_=28.3, p<0.0001) and interaction (F_(43,528)_=1.9, p<0.001). The sum of daily infrared beam breaks remained significantly higher in knockout animals (Fig. 2D, p<0.05, Student’s t-test). There were no differences in the alpha duration and period length between genotypes (data not shown). Interestingly the elevated locomotor activity in the knockout mice at around ZT18 correlates with the maximum expression of *Kcnk9* mRNA in wildtype animals, as seen in Fig. 1A.

**Figure 2.**
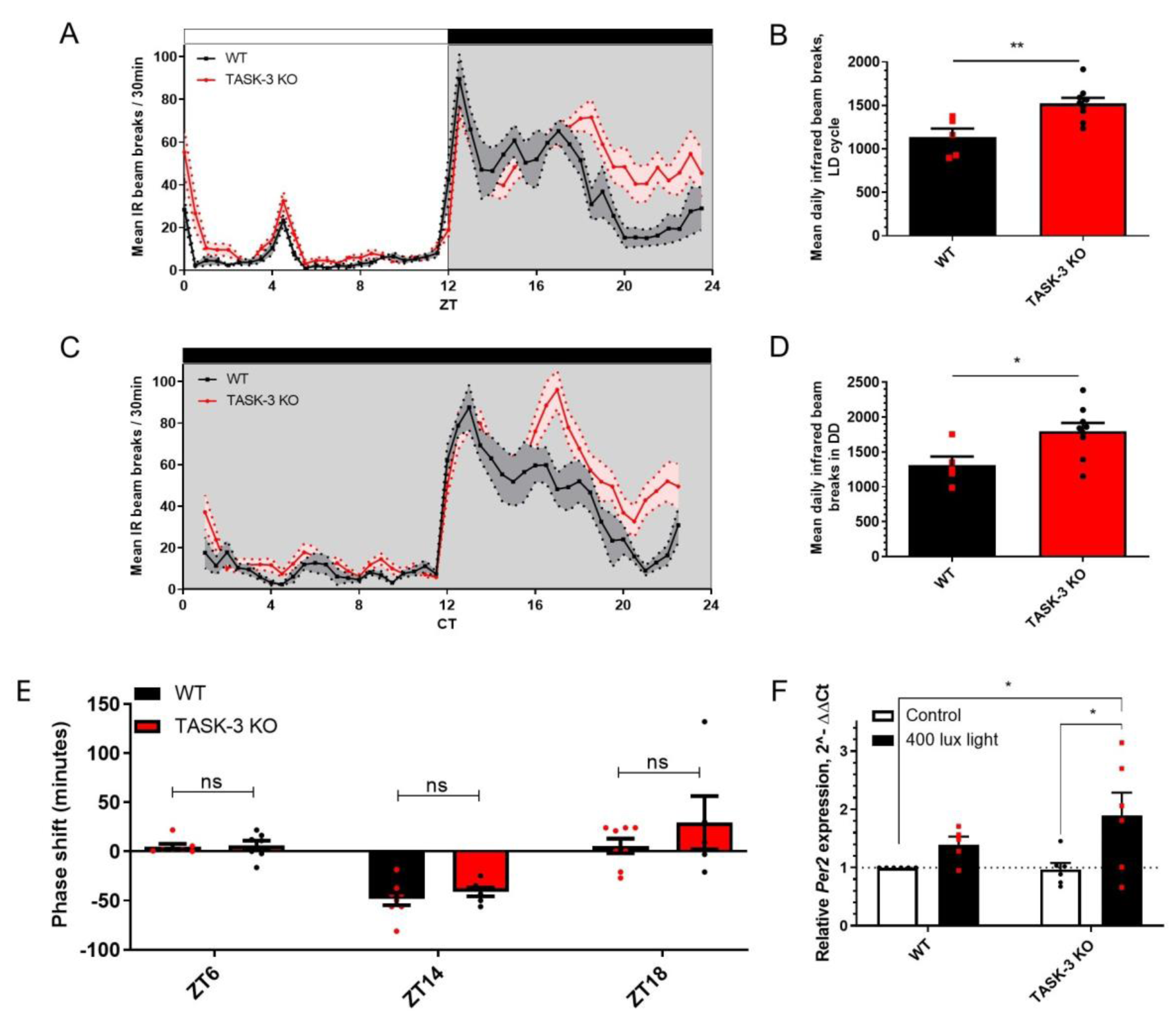
Locomotor activity and acute response to light in TASK-3 KO mice. (A-B) Infrared beam break recordings under 12:12 LD show higher TASK-3 KO activity. (C-D) Under constant dark housing, TASK-3 KO mice maintained significantly increased locomotor activity relative to WT. A-D: n=5 WT, 9 KO. (E) light pulses (400lux) presented in the middle of the day (10 min; ZT6,, n=7 WT/KO) or early/late night (5 min; ZT14; n=8 WT, 6 KO; and ZT18; n=8 WT, 4 KO) resulted in phase delays with no differences observed between genotypes. (F) Light induced Per2 expression was observed in both WT and TASK-3 KO mice, with 40% and 96% increases respectively (Tukey’s multiple comparisons test, n=5/6). *p<0.05, **p<0.01. Data presented as mean ± SEM.

### TASK-3 is not required for acute SCN responses to light

Brief light pulses during the middle of the day, early and late night (ZT6, ZT14 and ZT18 respectively) were used to assess the impact of TASK-3 loss in circadian behavioral resetting. A 400 lux light pulse, the same as ambient housing light intensity, induced a phase shift at ZT14 and 18 but this was not different between genotypes (Fig. 2E, two-way ANOVA with Tukey’s multiple comparisons test; genotype F_(1,35)_=1.8, p=0.2, interaction F_(2,35)_=0.7, p=0.5, time of day F_(2,35)_=23.4, p<0.0001). To confirm that the molecular response of the SCN to light was also still intact, we investigated the expression of *Per2* in response to light (Fig. 2F). There were no significant differences between genotypes. Two-way ANOVA analysis showed that significant source of variation was from light only (F_(1,19)_=9, p<0.01; genotype F_(1, 19)_=1.1, p>0.05; interaction F_(1, 19)_=1.4, p=0.2).

### Loss of TASK-3 dampens SCN’s endogenous cellular electrical firing rhythm and consequential responses to glutamate

We examined spontaneous SCN firing activity using acute SCN slices on multi-electrode arrays. Mean firing rate (MFR) measurements taken from SCN cells during the day reveal significantly dampened day time firing rate in TASK-3 KO’s compared to WT (p<0.0001, two-way ANOVA with Tukey’s multiple comparisons test, Fig. 3A). The source of variation is significant for genotype (F_(1,1762)_ = 40.2, p<0.0001), time of the day (F_(1,1762)_ = 98.3, p<0.0001) and interaction (F_(1,1762)_ = 29.9, p<0.0001, two-way ANOVA). The SCN in TASK-3 KO’s is silenced to WT night levels with respect to MFR. However, the difference between day and night-time firing rates remains significant (p=0.02). In order to ascertain if light transduction mechanisms were affected in TASK-3 KO mice, we applied glutamate, the neurotransmitter responsible for relaying light input from the retinohypothalamic tract to the SCN, to acute SCN slices. A concentration dependent increase in MFR in WT tissue samples was readily distinguishable (two-way ANOVA Tukey’s multiple comparisons test, control versus 30µM and 100µM glutamate concentrations p<0.0001, Fig. 3B). In contrast, SCN cells from TASK-3 KO mice did not exhibit such concentration-response effect, with quite consistent MFR responses, nevertheless significant only at 30 µM glutamate (p<0.05, two-way ANOVA Tukey’s multiple comparisons test, Fig. 3C). As we had previously observed (Lalic et al., in press), we noted that glutamate application resulted in heterogenous neuronal activity responses with some electrodes showing increased activity and other showing decreased activity upon adding glutamate. In wildtype animals 58% of electrodes showed an increase in activity, defined as an 10% or more increase in MFR upon addition of glutamate (Fig. 3D). However, in the TASK-3 KO animals 74% of electrodes responded by an increasing activity in response to glutamate (Fig. 3E). Overall, TASK-3 KO slices show a greater proportion of electrodes with increased activity after glutamate and a lower proportion with reduced activity compared to WT animals (Chi-square = 46.83; p<0.0001), suggesting a significant change in the excitatory-inhibitory balance in the SCN.

**Figure 3.**
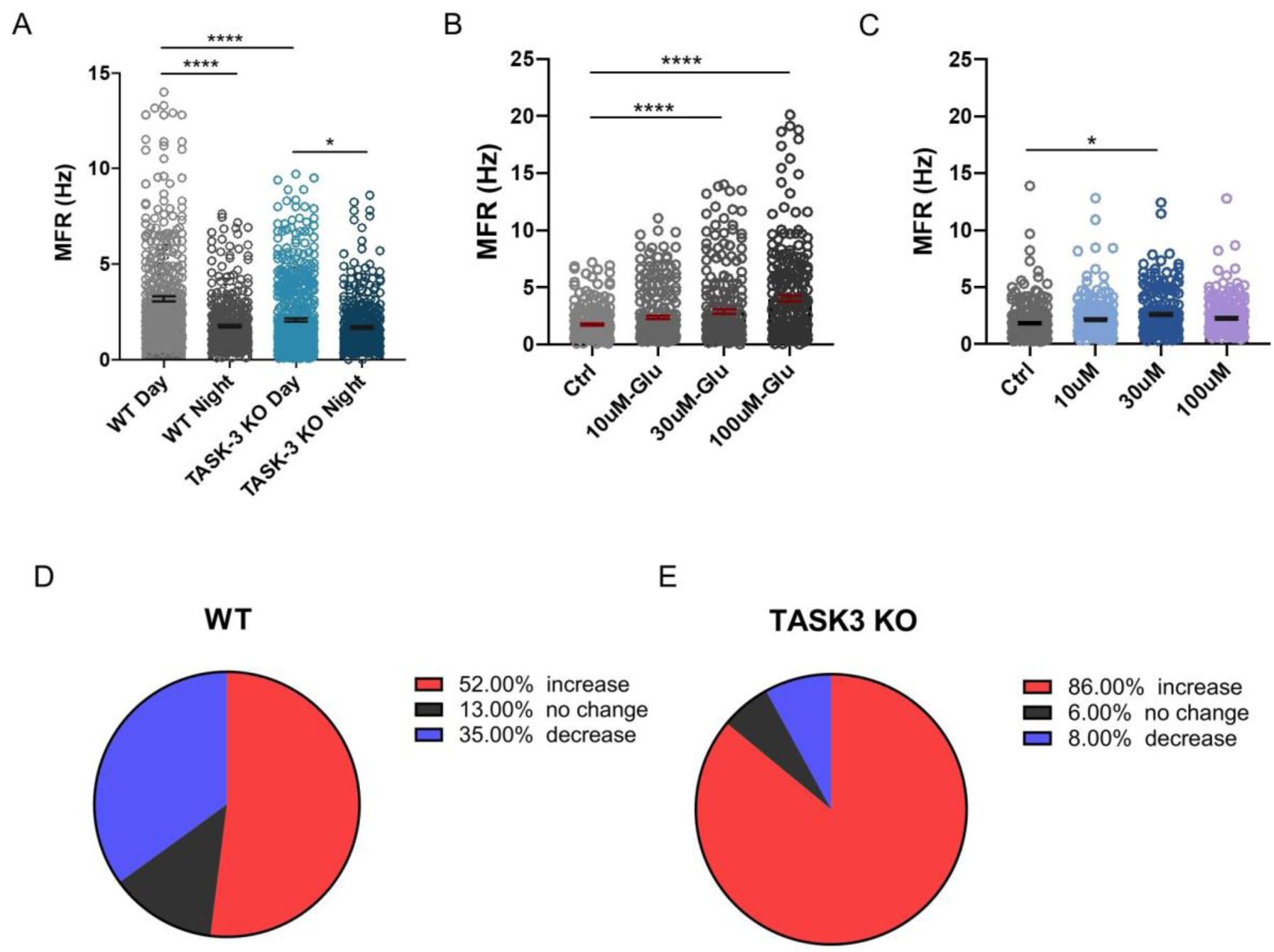
The role of TASK-3 in SCN firing. (A) Mean firing rate (MFR) in WT SCN shows diurnal variation, with significantly higher MFR levels during the day. This was significantly dampened to WT night levels in TASK-3 KO mice. MFR calculated by pooling data from all SCN electrodes (WT; day, n=7; night, n=8 animals; TASK-3 KO; day, n=5; night n=5 animals). (B-C) In vitro glutamate application on acute SCN slices shows a dose dependent increase in the mean firing rate in WT mice (B), but not in the TASK-3 Kos (C). (D-E) Categorizing activity of SCN neurons in response to glutamate, into those that showed reduced firing (>10%), no change in activity (+/− 10%) and increased firing (>10%), in comparison to basal and 100 μM glutamate. TASK-3 KO show a greater proportion of electrodes with increased activity after glutamate and a lower proportion with reduced activity (Chi-square=46.83; p<0.0001). All grouped data are mean ± SEM. *p<0.05, **p<0.01, ***p<0.001; ****p<0.0001.

### TASK-3 is required for rapid re-entrainment to ‘jet lag’ like phase shifts

Travel across a number of time zones in rapid succession requires significant adaptation by the SCN and leads to the phenomenon of ‘jet lag’. The mechanisms involved are poorly understood and typically, symptoms are short lived due to the rapid synchronization of the circadian system to the new day-night cycle. Given there were alterations in behavioral resetting responses to light in the absence of TASK-3, we set out to test the extent to which TASK-3 was involved in stable photo-entrainment of the circadian clock. We exposed our animals to a 6-hour advance in the light dark cycle to emulate a ‘jet lag’ scenario. On a daily basis, phase shifts differed between experimental groups, with WT animals having a greater daily advance, thus reaching entrainment to the new LD cycle much quicker. However, TASK-3 KO animals had a significantly slower rate of re-entrainment, with significant differences compared to WT from days 3 to day 11 (Fig. 4A, two-way ANOVA for genotype F_(1,207)_=157.7, p<0.0001, day after jet lag F_(15,207)_=89.6, p<0.0001, interaction F_(15; 207)_=2.6, p<0.01). The number of days needed to fully re-entrain to the advanced LD cycle was significantly increased in TASK-3 KO mice, taking 15 days compared with 8 days in WT controls (p<0.001, n = 8 WT, 7 TASK-3 KO, unpaired t-test). Interestingly, TRESK KO mice did not show any impairment in their rate of re-entrainment (Fig. S1), implicating TASK-3 as the key candidate for enabling SCN entrainment to challenging lighting environments.

**Figure 4.**
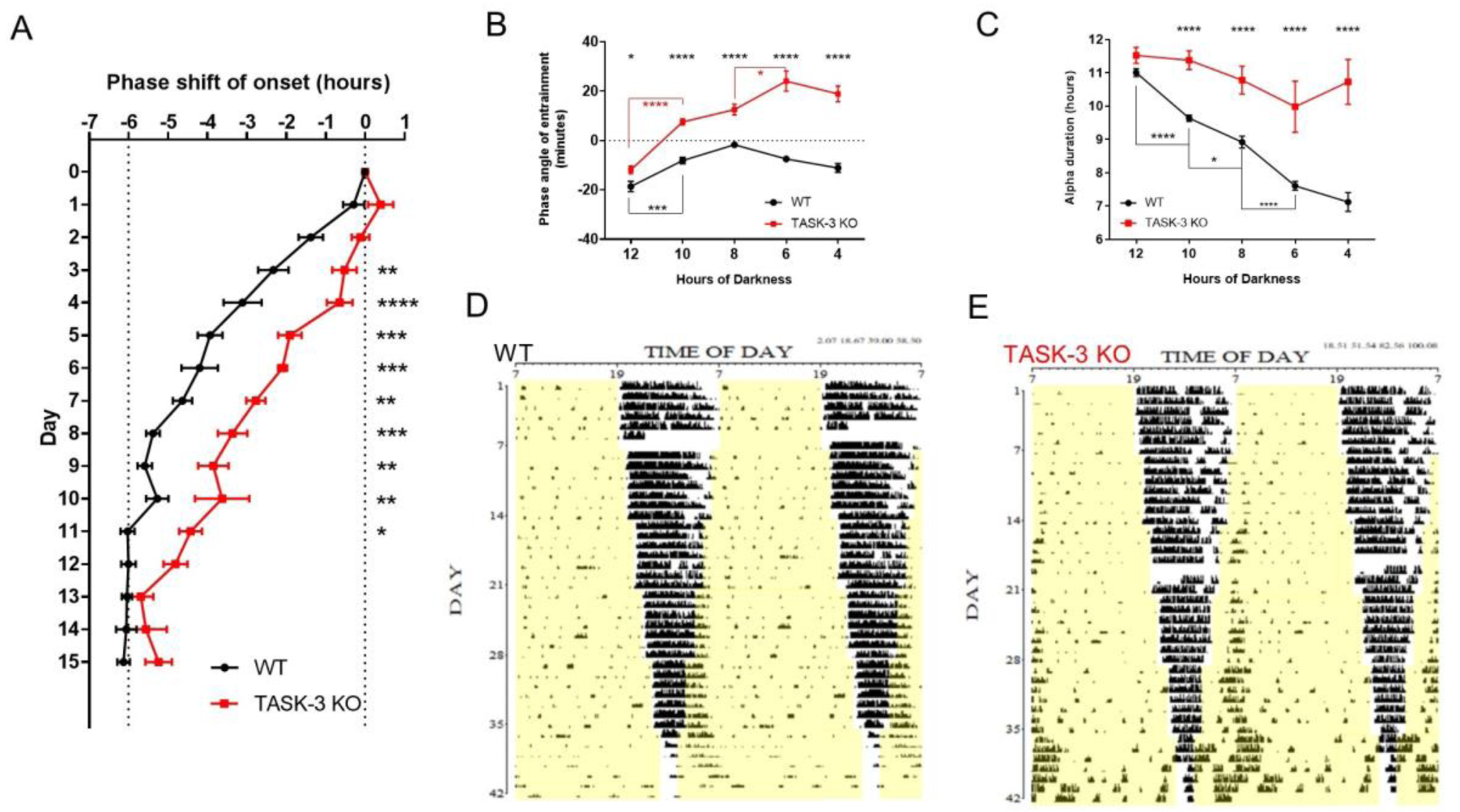
Behavioral response to challenging light conditions. (A) TASK-3 KO mice exhibit a reduced rate of re-entrainment to a 6-hour shift in the LD cycle. TASK-3 KOs take on average 4 additional days to reach stable entrainment (Tukey’s multiple comparisons test; n=8 WT, 7 KO). (B) TASK-3 KO mice fail to track activity onset to the compressed light-dark cycles, with WT animals maintaining stable phase angles of entrainment. Bonferroni’s multiple comparisons test showed significant differences between genotypes across all dark variations. (C) WT mice showed robust activity compression, predominantly restricted to the dark portion of the compressing light dark cycles. However, TASK-3 KOs were unable to compress activity in this manner. For figures B and C stars at the top of the graph show the significance between genotypes, whereas stars with connecting bars show significance within the genotype across consecutive data points (n=8 WT, 6 KO). (D-E) Representative actograms showing synchronization to compressed LD cycles in WT and TASK-3 KOs. *p<0.05, **p<0.01, ***p<0.001, ****p<0.0001. Data presented as mean ± SEM.

### Loss of TASK-3 results in impaired responses to progressively compressing light-dark cycles

Seasonal adaptation to changing durations in daylight length are key survival mechanism for many species. The ability to alter behavior following such variations in lighting levels is regulated by the SCN. We sought to test the contribution of TASK-3 in driving such adjustments. Animals were placed under gradually compressed LD cycles, with daylight hours lengthening every seven days, reaching constant light within six weeks. Wild type animals were able to track the ‘lights-off’ cue extremely well, with very little deviation from the start of this signal (Fig. 4B, 4D). However, TASK-3 KO animals exhibited significantly less robust entrainment (Fig. 4B, 4E; two-way ANOVA for genotype: F_(1,427)_=200.7, p<0.0001; the duration of darkness: F_(4,427)_=49.7, p<0.0001, interaction F_(4,427)_=13.9, p<0.0001). Further, TASK-3 KO mice showed a constant variation in entrainment as the daylight hours increased (one-way ANOVA, F_(4,195)_=39.4, p<0.0001; Fig. 4D).

WT mice were able to effectively compress most of their activity to fall within the dark portion of each compressing LD cycle. A one-way ANOVA for WT alpha duration during the increasing photoperiods showed significant differences in alpha (F_(4, 230)_=101.9, p<0.0001). Tukey’s multiple comparisons test showed a significant shortening of alpha with decreasing hours of darkness between 12-10 (****p<0.0001), 10-8 (*p<0.05) and 8-6 (****p<0.0001) hours of darkness. Lack of robust entrainment in TASK-3 KO mice continued throughout, with activity compression becoming increasingly impaired (Fig. 4C, 4E). A one-way ANOVA did not show significant differences between alpha durations through lengthening photoperiods in TASK-3 KO animals (F_(4, 144)_=2.077, p>0.05). Comparison between genotypes showed a significant difference across all compressed activity measurements (two-way ANOVA for genotype: F_(1, 374)_=120.4, p<0.0001, the hours of darkness: F_(4, 374)_=31.73, p<0.0001; interaction F_(4, 374)_=9.0, p<0.0001).

## Discussion

Our findings show that TASK-3, a K2P channel, plays a critical role in the plasticity of the circadian clock in driving behavioral synchronization to challenging environmental lighting conditions. From our work, TASK-3 is required for the circadian rhythm in both resting membrane potential and robust diurnal mean firing rates. Interestingly, it is the latter that appears to be the most significant, with the dampening of MFR in the absence of this channel rendering mice unable to rapidly synchronize to both shifted and compressed day-night cycles; a skill imperative for survival and higher-level wellbeing.

The ability to successively modulate the SCN clock for behavioral re-entrainment following shifts or durations in the light dark cycle is a fundamental function of the SCN. We found that TASK-3 KO mice took a greater number of days to re-entrain, thus the symptoms associated with jet lag would have persisted longer in these animals compared to their WT counterparts. Apart from re-entrainment when travelling across time zones, our bodies also need to adapt to the changing celestial cycles. By using a lengthening daylight, compressing night-time, paradigm we found that TASK-3 KO mice struggled to track lights off and compress their activity to the dark phase, as the duration of darkness reduced over time, emulating light changes during the seasonal summertime. This behavioral data provides strong evidence for the role of TASK-3 in providing the SCN with an element of neural plasticity that allows it to react and adapt to changing environmental light-dark cycles; a fundamental property of circadian synchronization.

Surprisingly, it would appear that TASK-3 is not involved in acute phase resetting to light pulses, with TASK-3 KO mice showing normal phase resetting in responses to advancing and delaying photic stimuli. This is despite acute responses to glutamate on acute SCN slices being dampened in TASK-3 KO compared to wildtype mice. The mechanism leading to impaired re-entrainment in TASK-3 knock-out mice is therefore unexplained. Our findings suggest a more subtle change in the intra-SCN communications and plasticity of SCN neurons. Certainly, the electrical properties of the SCN are fundamental to its ability to respond to challenging light environments. In this respect it is interesting that the resting membrane potential (RMP) in the TASK-3 KO mouse resides at the same level as that of the wild type day-time level. This would imply that the SCN in these animals remains in a constant day like state with regards to RMP. However, the link between resting membrane and excitability is often complex. Intuitively one might envisage that loss of a hyperpolarizing brake (in this case TASK-3 activity) might increase excitability and therefore firing rates. Indeed, we found that SCN neurons were more likely to respond to glutamate by increased firing but despite this, the spontaneous MFR was dampened. TASK-3 has previously been shown to have an additional, critical role in resetting the membrane potential between action potentials and so loss of TASK-3 can result in lower firing rates as we have observed (Brickley *et al.*, 2007).

Interestingly, our findings here are distinct from the work we have undertaken in TRESK knock-out mice, where we also observed a loss of diurnal variation in RMP (Lalic *et al*, in press) and a depolarized RMP both in the day and night. For the TRESK knock-out the MFR is elevated rather than dampened. This may relate to the role of TRESK in modulating the threshold potential (Pettingill *et al.*, 2019; Weir *et al.*, 2019), whilst TASK-3 may have a more dominant role in after-hyperpolarization. Furthermore, TRESK KO mice do not show altered circadian behavioral entrainment phenotypes which are present in TASK-3 KO animals. Thus, taken together, this would suggest that both TASK-3 and TRESK regulate MFR in the SCN, however, it is TASK-3 that is responsible for neural plasticity that drives circadian adaptations to environmental changes such as jet lag or seasonal variation in day-night durations. TRESK appears to play an important complementary role in acute clock resetting and acute glutamate responses, but is unable to drive clock synchronization to seasonal and dramatic changes in daylight durations; this role being reserved for TASK-3.

The dampened MFR in TASK-3 KO mice may lead to an overall reduced electrical coupling between internal SCN regions as well as efferent targets, thus impacting SCN communication. If this also interferes with ventral dorsal synchrony, in circumstances such as jet lag and daylight lengthening conditions, where the SCN is having to continuously re-synchronize to the external light–dark cycle may be compromised. We propose this impaired synchrony generates the defective behavioral phenotypes we observe here and will be the subject of future studies.

In conclusion, our work has demonstrated the important role played by the TASK-3 K2P channel in the regulation and maintenance of stable mammalian circadian rhythms. This channel provides the SCN with the ability to rapidly adapt to changing environmental daylight cycles and possibly controls the synchronization of the pacemaker. Finally, one might have expected acute phase responses to light to be the main driver for jet lag and seasonal changes, however our work shows that the mechanisms may be very distinct. More generally, TASK-3 is a vital component in the circadian synchronization of physiological and behavioral rhythms, thus is a key part in the overall health and wellbeing of an organism.

## Materials and Methods

### Animals

TASK-3 KO animals were gifted by William Wisden. The generation of TASK-3 mice can be found in Brickley et al. paper [4].

Males were used for all behavioral recordings and molecular work. Males and females were used for the electrophysiology recordings.

All behavioral testing was performed at Charles River Animal Facility using infrared sensors (LuNAR™ PIR 360°, Risco Goup) or 8 cm internal diameter cardboard running wheels. The data was collected using The Chronobiology Kit (Stanford Software systems). Activity counts per minute were recorded and data was saved to the computer every hour. All mice were bred on a C57BL/6J background from heterozygous parents generating TASK-3 KO and littermate wild type animals. Male mice were housed individually in polypropylene cages with food and water available ad libitum.

All procedures complied with the UK Animals (Scientific Procedures) Act (1986) and were performed under a UK Home Office Project License in accordance with University of Oxford and University of Kent Policy on the Use of Animals in Scientific Research. This study conforms to the ARRIVE guidelines.

### 24-hour clock gene expression

Age-matched male mice were sacrificed by cervical dislocation at ZT 2, 6, 10, 14, 18, 22 (3 mice per time point). During ZT 2-10 brains were removed under the room light, whereas during ZT 14-22 brains were removed under dim red light. To prevent SCN from signaling from the retina, eyes were immediately removed from animals sacrificed during ZT 14-22. 1mm coronal brain slices were made using brain matrix, and SCN was dissected under the stereomicroscope. Dissected SCNs were immediately submerged in RNAlater (Sigma, UK), frozen on dry ice and then kept at −80°C. RNA was isolated from tissue extracts using RNeasy Mini Kit (QIAGEN, UK). The RNA concentration was measured using Qubit RNA HS Assay Kit (ThermoFisher), and RNA was reverse transcribed by Nanoscript reverse transcription kit (Primerdesign, UK) with the final working cDNA concentration of 1ng/μL. 20μL reactions were prepared in triplicate in 96-well white plates (Alpha Laboratories, UK) consisting of 10μL iTaq™ Universal SYBR® Green Supermix (BioRad), 1μL 20x TaqMan gene expression assay, 6μL RNase-free water and 3μL cDNA (For TRESK expression 7μL of cDNA were used). Reactions were run using Applied Biosystems 7500 fast Real-Time PCR Systems.

20x TaqMan gene expression assays:

<u>Mm02014295_s1 Task-3</u>

<u>Mm02619580_g1 Beta-actin</u>

Relative expression was calculated using 2^−ΔCt^, where ΔCt is Ct (gene of interest) – Ct (reference gene – beta-actin)(Schmittgen and Livak, 2008). Linear detrending and normalization to mean of the 2^−ΔCt^ values was done using BioDare2 online tool. Data was standardized using the equation:

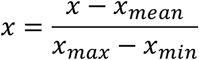

Graphs were plotted using GraphPad Prism 8 and standard sine wave was fitted to the data.

### Electrophysiology

For whole-cell patch-clamp studies 320μm coronal slices containing SCN were prepared in ice-cold high-sucrose aCSF containing 85 mM NaCl, 25 mM NaHCO3, 2.5 mM KCl, 1.25 mM NaH2PO4, 0.5 mM CaCl2, 7 mM MgCl2, 10 mM glucose, and 75 mM sucrose after decapitation under isoflurane anesthesia. Slices were then transferred to oxygenated aCSF (95% O2/5% CO2) containing 130 mM NaCl, 25 mM NaHCO3, 2.5 mM KCl, 1.25 mM NaH2PO4, 2 mM CaCl2, 1 mM MgCl2, and 10 mM glucose and then maintained at room temperature until recording.

During recordings, slices were superfused with aCSF saturated with 95% O2/5% CO2 at room temperature. Whole-cell patch-clamp electrodes (4–7 MΩ) were filled with an intracellular solution containing 120 mM K-gluconate, 10 mM KCl, 10 mM HEPES, 4 mM MgATP, 0.3 mM NaGTP, 10 mM Na-phosphocreatine, and 0.5% biocytin. Slices were viewed with an upright microscope (Slicescope -Olympus BX51, Scientifica, UK), using an air objective (2.5x) to visualize the whole slice and locate SCN followed by water immersion lens (40x) and IR-DIC optics. Cells were typically visualized from 30 to 100µm below the surface of the slice. The image was detected using a sensitive and fast camera (Photometrics Prime 95B sCMOS, Cairn, UK).

Resting membrane potential was measured immediately upon establishing whole-cell access. Resting membrane potential and spontaneous firing was recorded in bridge mode while zero current was inputted (I=0 mode). Intracellular recordings were obtained using a Multiclamp 700B amplifier and digitized at 10–20 kHz using Digidata 1440A acquisition board. Biocytin was included in the intracellular solution to allow post hoc visualization and confirmation of cell identity and anatomical region. All data were analyzed offline with Clampfit (pClamp 10), Neuromatic (http://neuromatic.thinkrandom.com), and custom-written software running within IgorPro environment.

### Locomotor activity during LD and DD housing

5 WT and 9 TASK-3 KO males were housed under 12:12 LD cycle and locomotor activity was recorder IR sensors. Environmental lighting was administered using LED strips fitted above each row of cages providing equal irradiance to all animals. LD cycle was controlled by a programmed timer, and light intensity was adjusted using neutral density filters. 24-hour LD activity profile was plotted by extracting data in 30 min bins over 9-10 days of activity recording (days, when cages were cleaned, were excluded). Mean 24-hour activity for each animal was calculated and plotted (mean ± SEM).

Free-running behavioral rhythms were measured from animals housed in 10 days of constant darkness following stable entrainment to a LD cycle. Data was extracted in 10 min bins for daily IR beam breaks or in 30 min bins for 22-hour behavior profile.

Activity profiles were determined by ‘eye-fitting’ regression lines to activity onsets under DD. Each daily intersection along this line determined the activity onset, thus CT 12. From CT 12, −11/+11 hours of activity data were used to plot the 22-hour profile.

### Circadian clock resetting with a light pulse

The phase shifting experiments were done using Aschoff type II protocol. 8 WT and 6 TASK-3 KO mice (4 during ZT18 phase shifting) were entrained to 12:12 LD cycle with 400 lux illumination during the light period. After stable entrainment, for at least seven days all mice received a 5min 400 lux light pulse at ZT14. Experiment was also done at ZT18. For the daytime (ZT6) light pulse 7 WT and 7 TASK-3 KO animals were entrained to 12:12 LD cycle with 400 lux illumination during the light period. On the day of experiment, lights did not turn on and 10min light pulse was administered at ZT6.

All mice were in the home cages during light pulse administration and were not moved to a new location. After the light pulse mice were left to free-run in constant darkness for 9-10 days. Cages were not cleaned two days before the light pulse and throughout the free-running phase. Light pulse induced clock gene changes

5/6 mice male mice per group were housed under the 12:12 LD cycle with 400 lux light intensity during the light period of the day. On the day of experiment, controlled mice were removed from the room and experimental animals received 5min 400 lux light pulse at ZT14. SCN was removed at ZT15 for all the mice. SCN removal was done under dim red light. Dissected SCNs were snap frozen on dry ice and then kept at −80°C.

cDNA preparation was done as described previously. For the RT-qPCR, 20μL reactions were prepared in triplicate in 96-well white plates consisting of 10μL PrecisionPLUS qPCR Master Mix with SYBR green (Primerdesign, UK), 5.2μL nuclease-free water, 0.9μL 10μM of each forward and reverse primers and 3μL cDNA. Relative expression was calculated using 2^−ΔΔC^t method, and all values were normalized to WT control.

Primers:

Beta-actin

F: 5’ACCAACTGGGACGATATGGAGAAGA-3’

R: 5’CGCACGATTTCCCTCTCAGC-3’

Per2

F: 5’GTCCACCTCCCTGCAGACAA-3’

R: 5’TCATTAGCCTTCACCTGCTTCAC-3’

### Extracellular action potential recordings (MEA)

The SCN slices were placed on MEA array with 256 electrodes arranged in a 16×16 grid with inter-electrode distance of 100µm. Signals from all electrodes on the 256 MEA array were collected simultaneously using USB-256MEA channel system (Multichannel Systems, Germany). Two minute recordings were collected and repeated three times at 20kHz. Channels within the SCN were identified visually, and spontaneous extracellular action potentials recordings from these SCN channels were discriminated offline using threshold-based event counting in custom written scripts in IgorPro. Thresholds were typically set at 3.5x the baseline noise level, with typical signal amplitude between 20 and 75µV. Single unit discrimination was not routinely possible in these experiments. Custom built MATLAB software was used to compare changes of activity at highest dose of glutamate to baseline. If more than 10% increase found, these neurons were grouped in the increased group. If more than 10% decrease found, these were grouped in the decreased group. No change group contains neurons with 10% fluctuation above or below the baseline. Significance analyzed with one-way ANOVA test.

### Statistical Analysis

Data are represented as means ± SEM, and “n” refers to the number of observations. The number of animals in each data set is ≥3. Comparisons for statistical significance were assessed by one- or two-way ANOVA and post hoc multiple-comparison t tests or unpaired t tests using GraphPad Prism 8.

### Drugs

L-Glutamic acid was purchased from Tocris Bioscience. Drugs were added to oxygenated aCSF to final concentration immediately before use.

To assess the effect of glutamate on MEA activity of acute SCN slices, we bath applied different concentrations of glutamate (0, 10, 30 and 100 µM) with recording commencing 5 minutes after starting perfusion and continuing for 2 minutes. The same slice was used with successively increasing concentrations of glutamate.

### Jet lag and seasonal compression

8 WT and 7 KO male mice were housed individually in cages equipped with 8 cm internal diameter cardboard running wheels. First, mice were entrained to the 12:12 LD cycle for 13 days. On 14th day, timer was advanced by 6-hour at ZT10. Animals were maintained under the new LD cycle until full re-entrainment. Re-entrainment of individual mouse was considered complete when total shift in activity onset (for advancing) or activity offset (for delaying) reached 6 hours ± 0.5.

During increasing photoperiod experiment, 8 WT and 6 KO mice were maintained for 13 days under 12:12 h LD cycle for collection of stable entrainment data. On day 14 the photoperiod was increased by 1 hour at dawn and dusk thereby providing 14:10h LD cycle which was maintained for 7 days. A 2-hour increase in light hours was performed weekly to provide 7 day periods of 16:8h, 18:6h, 20:4h, 22:2h LD cycles, with a final 2 week period in constant light, LL. Phase angle of entrainment measured over the first 13 days under 12:12h LD and then from day 3-5 following change in photoperiod with days 1 and 2 disregarded. Total active period, α, was also measured daily for the first 13 days under 12:12h LD and then from day 3-5 following change in photoperiod with days 1 and 2 disregarded. Alpha (α), was calculated by subtracting time of activity onset from the activity offset for a given day.

## Acknowledgments

We would like to thank Charles River (UK) for the daily care of animals. This study has received support from the Innovative Medicines Initiative Joint Undertaking under grant agreement n°115439, resources of which are composed of financial contribution from the European Union’s Seventh Framework Programme (FP7/2007-2013) and the Royal Society (RG100842).

---

**Fig. S1.**
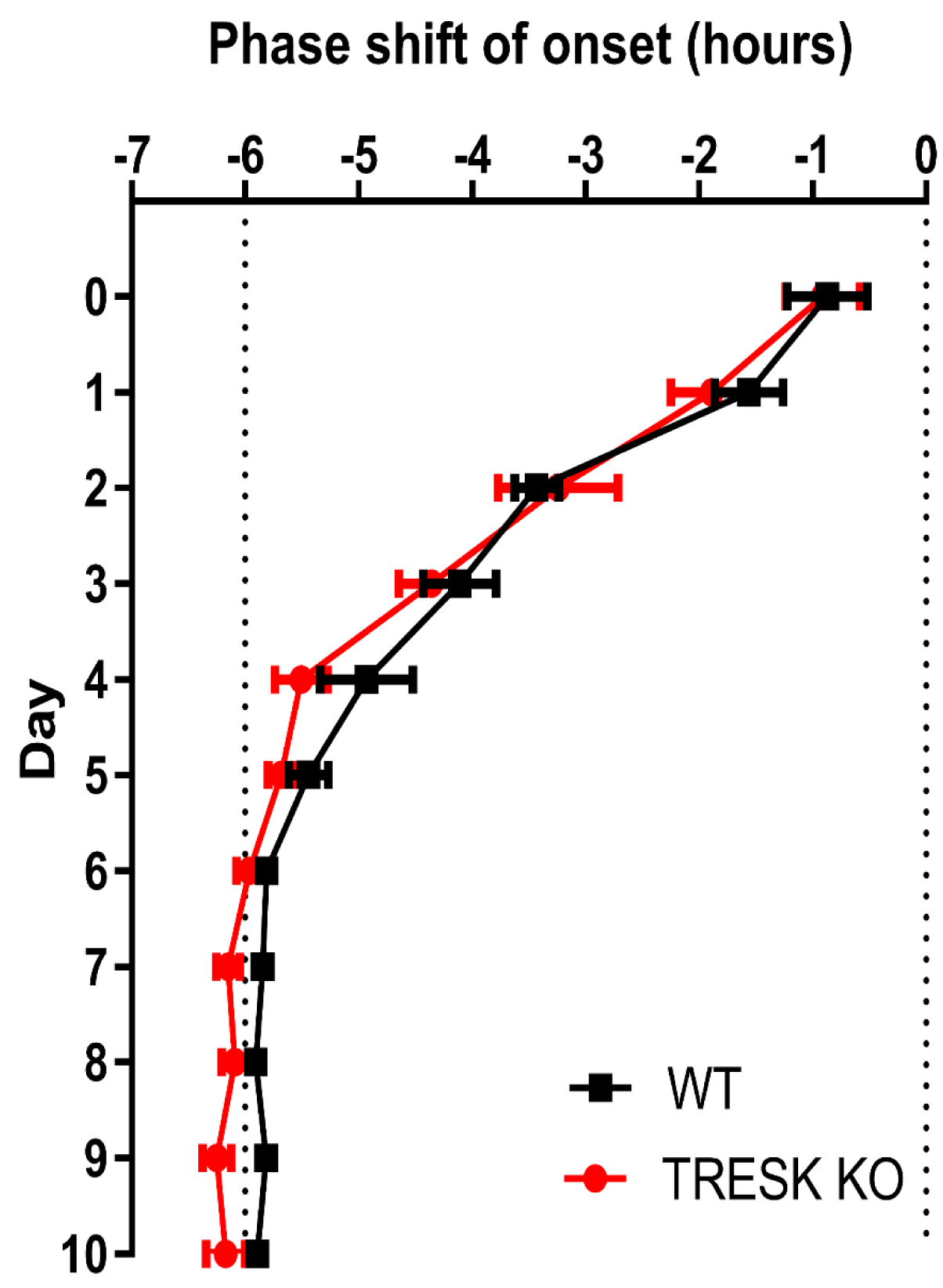
TRESK role in chronic resetting to light. SCN resetting to the 6-hour light-dark advance was nearly identical in WT and TRESK KO mice. n=5 WT, 5 KO; data presented as mean ± SEM.

